# Genetic screens identified dual roles of MAST kinase and CREB within a single thermosensory neuron in the regulation of *C. elegans* thermotaxis behavior

**DOI:** 10.1101/2022.07.12.499830

**Authors:** Shunji Nakano, Airi Nakayama, Hiroo Kuroyanagi, Riku Yamashiro, Yuki Tsukada, Ikue Mori

**Author notes:** Corresponding authors (S.N.), (I.M.). These authors contributed equally to this work (Airi Nakayama, Hiroo Kuroyanagi, Riku Yamashiro).

## Abstract

Animals integrate sensory stimuli presented at the past and present, assess the changes in their surroundings and navigate themselves toward preferred environment. Identifying the molecular and circuit mechanisms of such sensory integration is pivotal to understand how the nervous system generates perception and behavior. Previous studies on thermotaxis behavior of *Caenorhabditis elegans* suggested that a single thermosensory neuron AFD plays an essential role in integrating the past and present temperature information and is essential for the neural computation that drives the animal toward the preferred temperature region. However, the molecular mechanisms by which AFD executes this neural function remained elusive. Here we report multiple forward genetic screens to identify genes required for thermotaxis. We reveal that *kin-4*, which encodes the *C. elegans* homolog of MAST kinase, plays dual roles in thermotaxis and can promote both cryophilic and thermophilic drives. We also uncover that a thermophilic defect of mutants for *mec-2*, which encodes a *C. elegans* homolog of stomatin, can be suppressed by a loss-of-function mutation in the gene *crh-1*, encoding a *C. elegans* homolog CREB transcription factor. Calcium imaging analysis from freely-moving animals suggest that *mec-2* and *crh-1* function in AFD and regulate the neuronal activity of its post-synaptic interneuron AIY. Our results suggest that a stomatin family protein can control the dynamics of neural circuitry through the transcriptional regulation within a sensory neuron.

## Introduction

Information processing in the nervous system is essential for animals to survive and reproduce in response to changes in their environments. Research in the past decades have identified basic principles of the neural circuit operation that enable several functions of neural computations such as gain control of sensory stimuli and integration of multisensory information (Dunn and Rieke 2006; van Atteveldt *et al*. 2014). Identifying the site of such neural computations and deciphering the molecular and circuit mechanisms thereof are critical steps toward understanding how the nervous system generates perception and behavior.

Multisensory integration allows the animals to process stimuli of different modalities or of the same modality presented at different time points, and underlies decision making in the nervous system. A well-studied example includes the mushroom body, the learning center of the fly brain required for olfactory learning. Responses of the fruit fly to certain odors can be modulated by pairing the odorants with appetitive or aversive stimuli (Quinn *et al*. 1974; Tempel *et al*. 1983). The mushroom body receives inputs from both olfactory (Aso *et al*. 2014) and gustatory systems (Kim *et al*. 2017), suggesting that it is the site of multisensory integration that mediates olfactory learning. Studies in the fruit fly as well as in other species have thus elucidated a common neural basis for multisensory integration: multiple sensory inputs converge onto a single group of target neurons that integrate the signals and control the outputs.

Recent studies in the nematode *Caenorhabditis elegans* suggested a distinct circuit operation for integration of thermal stimuli presented at different time points. The wild-type animals that have been cultivated at a certain temperature with food migrate toward that cultivation temperature when placed on a thermal gradient (Hedgecock and Russell 1975). The *C. elegans* nervous system apparently integrates the past and the present temperature information and executes the appropriate behavior that drives themselves toward the cultivation temperature (Luo *et al*. 2014; Ikeda *et al*. 2020). Neural circuitry required for thermotaxis has been extensively studied (Mori and Ohshima 1995; Kuhara *et al*. 2008; Beverly *et al*. 2011; Ikeda *et al*. 2020). Central to this circuitry is the thermosensory neurons AFD and its postsynaptic interneurons AIY (White *et al*. 1986; Cook *et al*. 2019). The AFD neurons respond to temperature stimuli and increase intracellular calcium (Ca^2+^) level upon warming (Kimura *et al*. 2004; Clark *et al*. 2006, 2007; Ramot *et al*. 2008; Tsukada *et al*. 2016; Takeishi *et al*. 2016). The increase in the Ca^2+^ level of the AFD neurons reflects the information of the previous cultivation temperature and occurs within a temperature range with its lower bound determined by the previous cultivation temperature. Analysis of a primary cultured AFD neurons indicated that the temperature range of the AFD response is an intrinsic property of AFD and does not require the connection to the neural circuits (Kobayashi *et al*. 2016). Recent studies also suggested that in addition to these temperature-evoked Ca^2+^ responses, the AFD neuron modulates its neuronal outputs such that it evokes distinct responses in its post-synaptic neuron AIY (Hawk *et al*. 2018; Nakano *et al*. 2020). Thus, the single AFD neurons integrate the past and present temperature information and execute single-cell multisensory computation to achieve thermotaxis behavior. A similar neural operation has been also reported in the circuitry required for salt chemotaxis in *C. elegans* (Sato *et al*. 2021; Hiroki *et al*. 2022), suggesting that a single-cell integration is prevalent in the *C. elegans* nervous system. However, the molecular mechanisms by which the AFD neurons execute this modulation remain elusive.

To further understand the molecular basis of the single-cell computation during *C. elegans* thermotaxis, we here conducted forward genetic screens for mutants defective in thermotaxis. We show that *kin-4*, which encodes the *C. elegans* homolog of MAST (Microtubule-associated serine threonine) kinase (Walden and Cowan 1993), plays dual roles in thermotaxis and suggest that KIN-4 is a critical regulator of the single-cell computation within AFD. Our genetic screen also indicated that a thermotaxis defect of a stomatin homolog, *mec-2* (Huang *et al*. 1995 p. 2; Nakano *et al*. 2020), was suppressed by a loss-of-mutation in *crh-1*, which encodes the *C. elegans* homolog of CREB transcription factor (Kimura *et al*. 2002). Our results suggest that a stomatin family protein controls the multisensory integration in AFD via transcriptional regulation.

## Materials and Methods

### *C. elegans* strains

The *C. elegans* strains were cultured on NGM plates with the OP50 *Escherichia coli* as food (Brenner 1974). All strains were cultured at 20 °C unless otherwise indicated. N2 (Bristol) was used as the wild-type strain. Germline transformation was performed by microinjection as previously described (Mello *et al*. 1991). CRISPR-Cas9-mediated genome editing was performed as previously described (Dickinson *et al*. 2013; Dokshin *et al*. 2018). Mutations, extrachromosomal arrays, integrated transgenes used in this study were described in FileS1.

### Thermotaxis assay

Thermotaxis assays were performed as previously described (Ito *et al*. 2006). Two hermaphrodite animals at the fourth larval stage were placed onto a NGM plate and were allowed to lay eggs. Their F_1_ progeny from two NGM plates were collected, were washed with M9 buffer and were transferred onto the center of a thermotaxis assay plate that had been placed onto a temperature gradient from 17 °C to 23 °C with the gradient steepness of 0.5 °C/cm. The animals were allowed to freely move on the temperature gradient for one hour. The assay plate was divided into eight sections along the temperature gradient. The number of animals in each section was counted.

### Genetic screens for mutants defective in thermotaxis behavior

Wild-type animals were mutagenized by ethyl methansulfonate (EMS). Their F_2_ progeny were cultivated at 17 °C or 23 °C and were subjected to thermotaxis assays on a temperature gradient from 17 °C to 23 °C. Animals that had migrated to the 17 °C region when cultivated at 23 °C or to the 23 °C region when cultivated at 17 °C were picked as mutant candidates and were recovered onto NGM plates. We allowed each candidate animal to lay eggs and retested eight to twelve lines from each F_2_ candidate for thermotaxis behaviors.

To screen for mutations that can suppress the thermophilic phenotype of *mec-2(nj89gf)*, we mutagenized *mec-2(nj89gf)* animals with EMS, and their F_2_ progeny cultivated at 20 °C were subjected to thermotaxis assay on a temperature gradient from 17 °C to 23 °C. Animals that had migrated to the 17 °C region were picked as mutant candidates, and their progeny were retested for the suppression of the *mec-2(nj89gf)* thermophilic phenotype.

### Calcium imaging of the AFD neurons in immobilized animals

We generated animals expressing the calcium indicator YCX1.6 (Madisen *et al*. 2015) in the AFD and AIY neurons. The YCX1.6 in AFD was localized to the nucleus to separate the signals from AFD and AIY. The animals were immobilized by placing on a 10 % agarose pad with polystyrene beads (Polysciences), which were then covered by a cover slip. The samples were placed on a Peltier device used for the temperature control, and the YFP and CFP images were captured using epi-fluorescent microscope equipped with SOLA light engine (Lumencore) as a light source and were recorded at one frame per second with 400 msec exposure. Image processing was performed by MetaMorph software (Molecular Devices), and the fluorescent intensities of YFP and CFP were determined. The ratio change was calculated as (R_t_ - R_0_)/R_0_, where R_t_ represents the ratio of YFP to CFP of each frame, and R_0_ the mean ratio of the first ten frames.

### Calcium imaging of the AFD and AIY neurons in freely-moving animals

Animals expressing YCX1.6 in the AFD nucleus and the AIY neurons were placed on a 2 % agarose pad and were covered with a cover glass. The samples were placed on a motorized stage (HawkVision Inc.) with a transparent temperature-control device (TOKAI HIT Co. Ltd.). The animals were allowed to freely move on the agarose pad and were subjected to a temperature ramp. The YFP and CFP images were captured at 2 frames per second (30 msec exposure time) under epi-fluorescent microscope with SOLA light engine as a light source. The animals were kept under the field of view by controlling the stage movement, which was achieved by real-time analysis of transmitted infrared light images.

The image processing was first performed by DeepLabCut (Mathis *et al*. 2018; Nath *et al*. 2019) to extract the x-y coordinates of the region of the interest (ROI) for the fluorescent analysis of AFD and AIY. The image analysis was further performed by a custom-written program in MATLAB, and the positions of the ROI predicted by DeepLabCut were manually inspected for each frame. The AFD intensity was analyzed from its nucleus, and the AIY intensity was measured from a part of its neurite that makes a dorsal turn (White *et al*. 1986; Nakano *et al*. 2020). The ratio of fluorescence intensities (YFP/CFP) was used to calculate the standardized ratio change of AFD and AIY, which was defined as (R_t_ - R_min_)/(R_max_ - R_min_). The baseline standardized ratio, which was the mean of the standardized ratio values of the frames before the temperature was increased, was subtracted from the standardized ratio change of each frame. We calculated the area under the curve of the AIY standardized ratio change for the entire time window after the temperature stimulus was applied.

### Statistics

Normality of the data was assessed by Shapiro-Wilk test. Equal variance among data sets was assessed by Bartlett test. When both normality and equal variance were assumed, we used one-way analysis of variance (ANOVA) with Tukey-Kramer test or Dunnett test for multiple comparisons. Otherwise, we used Wilcoxon rank sum test.

## Results and Discussion

### A genetic screen for mutants defective in thermotaxis recovered 21 mutant isolates

To identify genes important for the regulation of thermotaxis, we conducted a genetic screen. We mutagenized the wild-type animals and looked for mutants that migrated toward the 23 °C region when cultivated at 17 °C or toward the 17 °C region when cultivated 23 °C (Fig. 1A). From this screen, we isolated 21 mutant strains, which were classified into three groups based on their thermotaxis phenotypes: nine mutants - *nj85, nj89, nj97, nj98, nj102, nj104, nj108, nj111* and *nj113* - displayed thermophilic phenotypes and migrated toward the higher temperature region (Fig. 1B); six mutants - *nj87, nj90, nj91, nj92, nj94* and *nj100* - showed athermotactic phenotypes and distributed evenly on the temperature gradient (Fig. 1C); and six mutants - *nj86, nj95, nj96, nj107, nj110* and *nj112* - exhibited cryophilic phenotypes and preferred the colder temperature region (Fig. 1D). We have previously reported that *nj89* is a gain-of-function allele of the gene *mec-2*, which encodes a *C. elegans* homolog of stomatin (Nakano *et al*. 2020) and that *nj90, nj94* and *nj100* are alleles of *kcc-3*, which encodes a potassium/chloride co-transporter that functions in a glial-like cell (Yoshida *et al*. 2016). In this study, we further characterize some of the thermophilic isolates, as described below.

**Figure 1.**
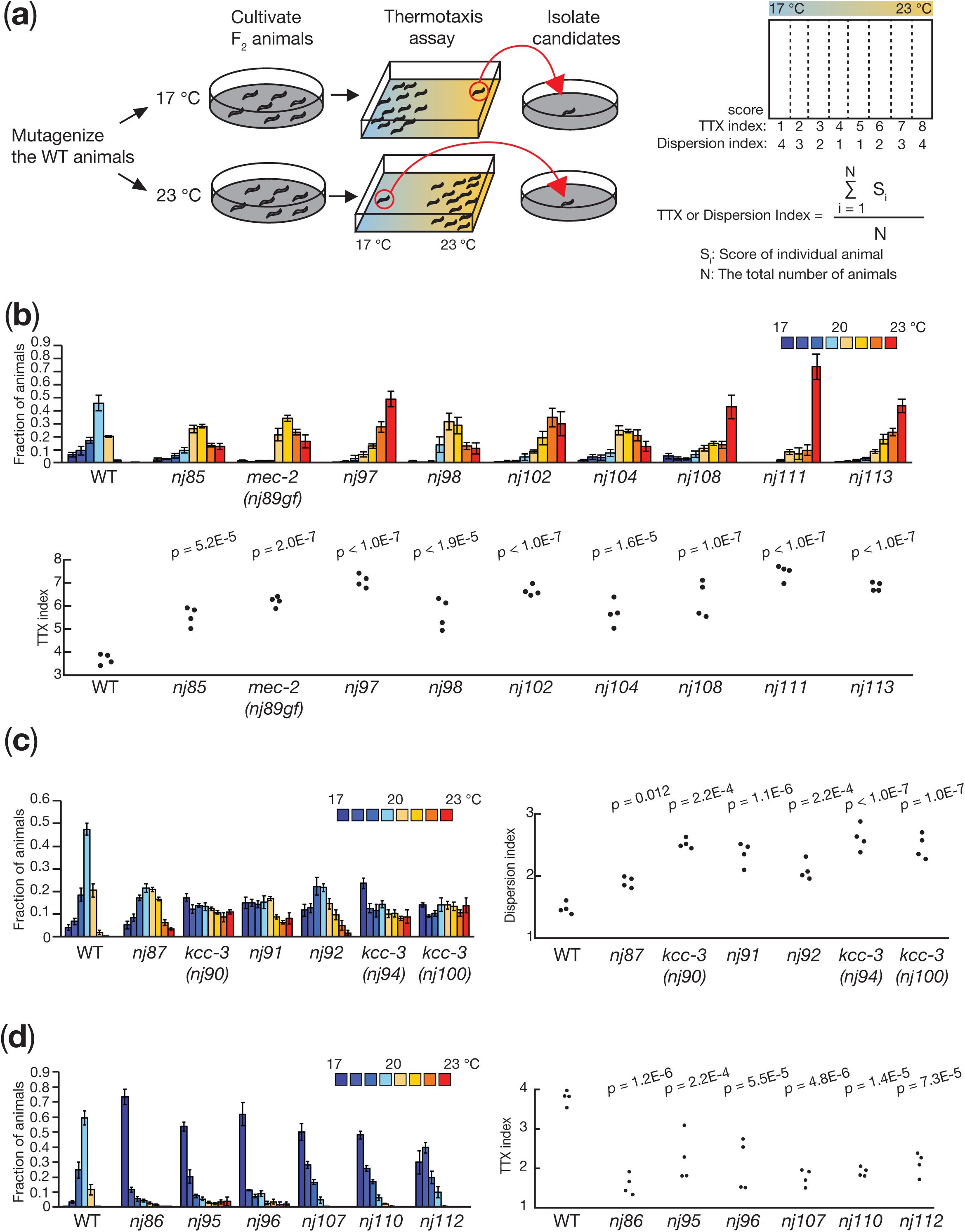
A forward genetic screen identified mutants defective in *C. elegans* thermotaxis. a) Schematic of the forward genetic screen and the formulas for TTX and Dispersion indices. We mutagenized the wild-type animals and screened their F_2_ progeny for mutants defective in thermotaxis. Animals that had migrated to the 23 °C or 17 °C region after cultivation at 17 °C or 23 °C, respectively, were isolated as mutant candidates. Thermotaxis behavior was quantified by counting the number of animals in each of the eight sections along the temperature gradient. We calculated TTX and Dispersion indices according to the formulas shown. b-d) Thermotaxis behaviors of thermophilic (b), athermotactic (c) and cryophilic (d) mutant isolates. Distributions of the animals in each section of the thermotaxis assay plate are shown as means ± SEM. TTX and Dispersion indices are shown as dots. *P* values were determined by one-way ANOVA with Tukey-Kramer test.

### *nj98* and *nj111* mutants carry mutations in the *pkc-1* locus

We observed that *nj98* and *nj111* failed to complement each other for their thermotaxis defects. To identify the gene responsible for the thermotaxis defects of these mutants, we mapped *nj111* into a 2.7 Mb interval on chromosome V (Figure 2). This region contains the *pkc-1* gene, which encodes a *C. elegans* homolog of protein kinase C-epsilon/eta. Our previous study indicated that *pkc-1*, also known as *ttx-4*, is required for thermotaxis (Okochi *et al*. 2005). We therefore asked whether *nj98* and *nj111* are alleles of *pkc-1*. We conducted DNA sequence analyses of these mutants and identified mutations in *pkc-1*: *nj98* carries a G-to-A transition mutation that is predicted to alter the glycine 1338 codon of *pkc-1c* to an aspartic acid codon; *nj111* is associated with a C-to-T transition mutation that would alter the arginine 80 codon to an opal stop codon (Figure 2). These observations suggested that *nj98* and *nj111* are alleles of *pkc-1*.

**Figure 2.**
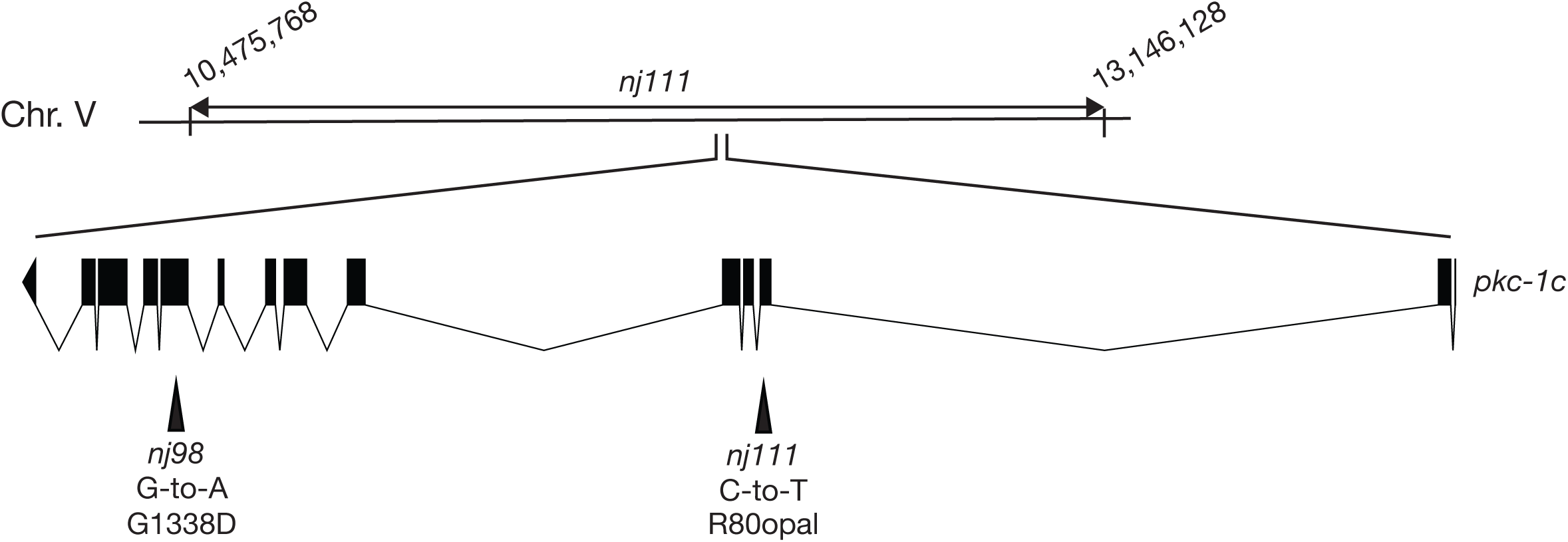
*nj98* and *nj111* carry mutations in the *pkc-1* locus. A chromosomal region to which *nj111* was mapped is indicated. The gene structure of *pkc-1c* is shown. The black boxes indicate exons, and the lines between the black boxes represent introns. Mutations identified in *nj98* and *nj111* are shown. The nucleotides correspond to the sequence in the sense strand.

### *nj97* is an allele of *pkc-2*

To identify the gene responsible for the thermophilic defect of *nj97* animals, we mapped *nj97* into a 1.9 Mb region of chromosome X. This region contains the gene *pkc-2*, which encodes a *C. elegans* homolog of protein kinase C beta (Fig. 3A). A previous study indicated that *pkc-2* is required for thermotaxis and that *pkc-2* functions in the AFD thermosensory neuron to regulate thermotaxis (Land and Rubin 2017). We therefore asked whether *nj97* is an allele of *pkc-2*. We identified a G-to-A transition mutation in the *pkc-2* locus of *nj97* animals that is predicted to alter the tryptophan 248 codon of *pkc-2a* to an amber stop codon (Fig. 3B). Introduction of a genomic clone that harbors the *pkc-2* locus rescued the thermophilic phenotype of *nj97* animals (Fig. 3C). These results indicated that *nj97* is an allele of *pkc-2*.

**Figure 3.**
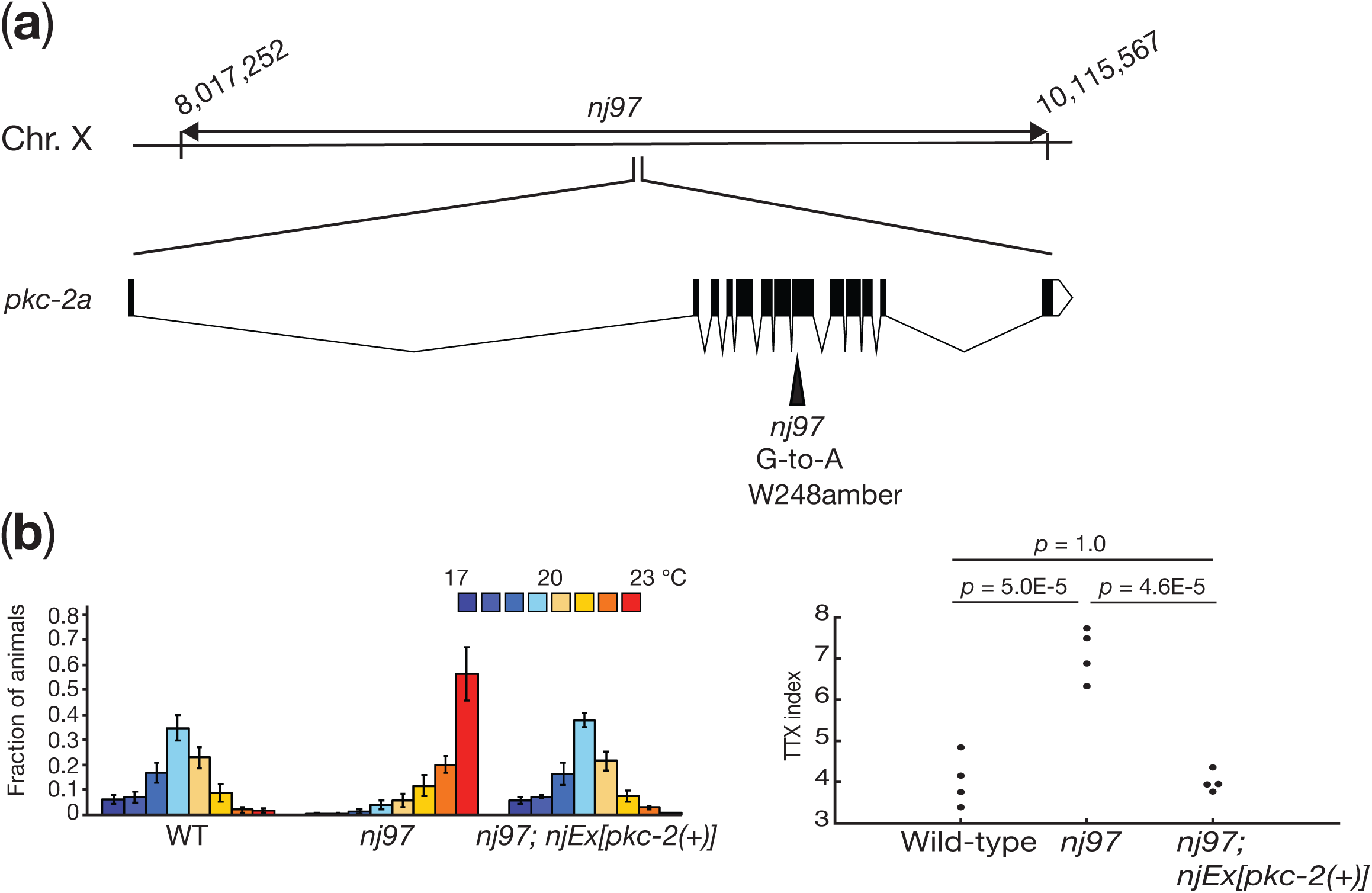
*nj97* is an allele of *pkc-2*. a) A chromosomal region to which *nj97* was mapped is indicated. The gene structure of *pkc-2a* is shown. The black boxes indicate exons, and the lines between black boxes represent introns. A mutation identified in *nj97* is shown. b) Thermotaxis behaviors of the wild-type, *nj97* and *nj97* carrying a genomic clone containing the *pkc-2* locus. Distributions of animals on the temperature gradient are indicated as means ± SEM. TTX indices are shown as dots. *P* values were determined by one-way ANOVA with Tukey-Kramer test.

### *nj104* and *nj108* harbor mutations in the *plc-1* locus

*nj104* and *nj108* failed to complement each other for the thermotaxis defect. To identify the gene responsible for their thermotaxis defects, we mapped *nj108* into a 180 kb interval of chromosome X (Fig. 3). This region contains the *plc-1* locus (Kunitomo *et al*. 2013), which encodes a *C. elegans* homolog of phospholipase C (PLC). PLCs cleaves phosphatidylinositol 4,5-bisphosphate (PIP_2_) into inositol 1,4,5-triphosplhate (IP_3_) and diacyl glycerol (DAG), the latter of which is known to act as a second messenger and can bind to and regulate diverse intracellular signaling proteins, including protein kinase C (Brose *et al*. 2004). We conducted DNA sequencing analyses of the *plc-1* locus in *nj104* and *nj108* animals and observed that these mutants carry mutations in *plc-1*: *nj104* is associated with a C-to-T transition mutation that is predicted to alter the glutamine 1112 codon of *plc-1d* to an ochre stop codon; *nj108* harbors a G-to-A transition mutation in the splice acceptor sequence within the 5th intron of *plc-1d*. These observations suggested that *nj104* and *nj108* are alleles of *plc-1*.

Our genetic screen identified three genes, *pkc-1, pkc-2* and *plc-1*, all of which were shown to be involved in the DAG signaling pathway: The PKC-1 and PKC-2 proteins contain the C1 and C2 domains that can bind to DAG (Corbalán-García and Gómez-Fernández 2014), and the PLC-1 protein promotes the production of DAG. Previous studies focusing on the *C. elegans* salt chemotaxis also indicated the importance of the DAG signaling in the ASER chemosensory neurons (Ohno *et al*. 2017). These observations suggest that the DAG signaling plays an important role in the single-cell computation within the *C. elegans* sensory neurons.

### *nj102* is an allele of *kin-4*

To identify the gene mutated in *nj102* animals, we mapped this mutation into a 300 kb interval of chromosome IV. This region contains the gene *kin-4*, which encodes the *C. elegans* homolog of microtubule-associated serine threonine (MAST) kinase (Fig. 5A). DNA sequencing analysis revealed that *nj102* animals carry a C-to-T transition mutation that alters the glutamine 1480 of *kin-4d* to an amber stop codon. We found that the introduction of a genomic clone carrying the *kin-4* locus rescued the thermophilic defect of *nj102* mutants (Fig. 5B). These results indicated that *nj102* is an allele of *kin-4*.

**Figure 4.**
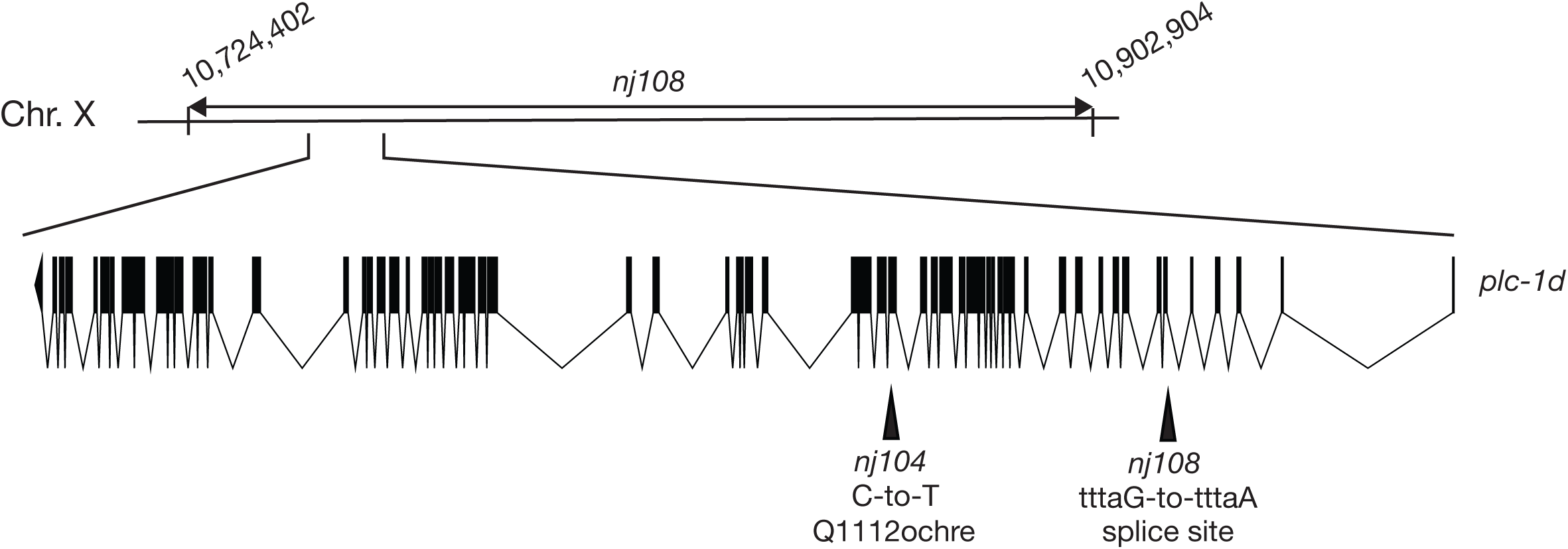
*nj104* and *nj108* harbor mutations in the *plc-1* locus. A chromosomal region to which *nj108* was mapped is indicated. The gene structure of *plc-1d* is shown. The black boxes indicated exons, and the lines between the black boxes represent introns. Mutations identified in *nj104* and *nj108* animals are shown.

**Figure 5.**
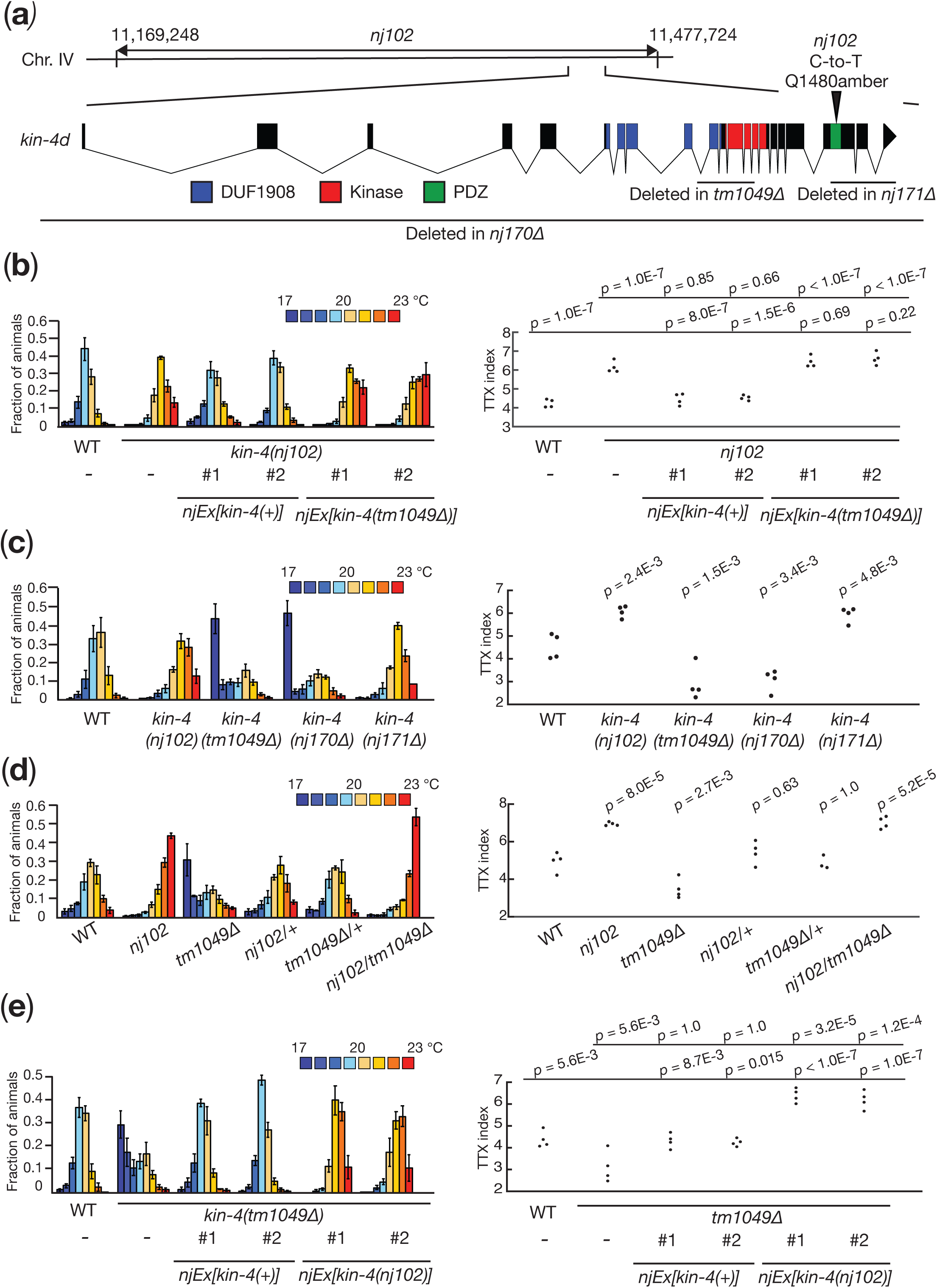
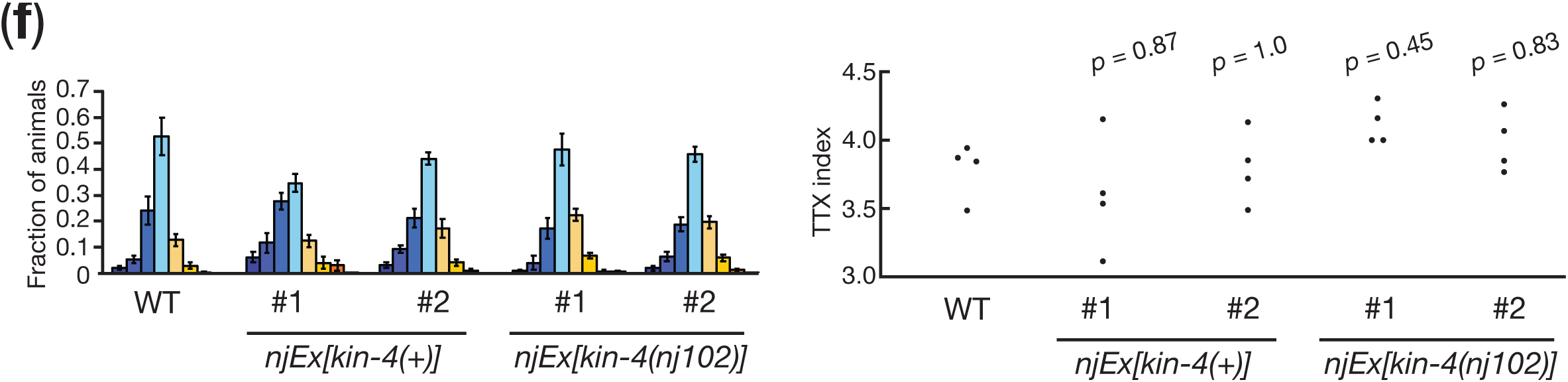
*nj102* is an allele of *kin-4*. a) A chromosomal region to which *nj102* was mapped is indicated. The gene structure of *kin-4d* is shown. The boxes indicate exons, and the lines represent introns. The colored boxes denote the coding sequences corresponding to the DUF1908, the kinase and the PDZ domains. Mutations associated with each mutant are shown. b)-f) Thermotaxis behaviors of *kin-4* mutants. Distributions on the temperature gradients are shown as means ± SEM. TTX indices are indicated as dots. *P* values were determined by one-way ANOVA with Tukey-Kramer tests in b), d), e) and f), and by one-way ANOVA with Dunnett test in c).

To further characterize the role of the *kin-4* gene in the regulation of thermotaxis, we analyzed other alleles of *kin-4*. As we previously reported (Nakano *et al*. 2020), two deletion alleles of *kin-4, tm1049* and *nj170*, displayed phenotypes distinct from that of *nj102* animals: *kin-4(tm1049)* and *kin-4(nj170)* mutants displayed bimodal distributions on the temperature gradient, with the majority of animals migrating toward the colder temperature region, while a minor population stayed around the cultivation temperature region (Fig. 5C). These results indicated that the majority of the *kin-4* null mutant animals display the cryophilic phenotype and that *kin-4* null phenotype is distinct from that of *kin-4(nj102)* animals.

The KIN-4 protein contains three major domains: DUF1908 (domain of unknown function), serine-threonine kinase domain and PDZ domain. *nj102* mutants harbor a mutation that would truncate the KIN-4 protein within the PDZ domain (Fig. 5A). To assess whether mutations that would eliminate the PDZ domain from the KIN-4 protein could cause the thermophilic phenotype similar to that of *kin-4(nj102)*, we generated another deletion allele of *kin-4, nj171. kin-4(nj171)* removes a part of the coding sequence of KIN-4 polypeptide that corresponds to the PDZ domain and its carboxy-terminal end (Fig. 5A). We found that *kin-4(nj171)* displayed a thermophilic phenotype similar to that of *kin-4(nj102)*. These results suggested that mutations that would eliminate the PDZ domain and C-terminal end of KIN-4 result in thermophilic phenotypes.

### *kin-4(nj102)* is likely a reduction-of-function mutation

To further characterize the nature of the *kin-4(nj102)* allele, we examined the thermotaxis phenotypes of a series of trans-heterozygotes as well as transgenic lines. First, both *nj102/+* and *tm1049/+* heterozygous animals showed the wild-type phenotype (Fig. 5D), indicating that *nj102* and *tm1049* cause recessive phenotypes. To ask whether *kin-4(nj102)* causes a gain-of-function mutation, we injected a genomic clone containing the *kin-4(+)* or *kin-4(nj102)* gene into the wild-type animals. Neither clone affected the thermotaxis phenotype (Fig. 5E), suggesting that *nj102* is not a gain-of-function mutation. When we examined the trans-heterozygotes of *nj102/tm1049*, these animals displayed a thermophilic phenotype similar to that of *nj102* animals (Fig. 5D). These observations suggested that *nj102* might be a reduction-of-function mutation. We therefore asked whether introduction of the *kin-4(nj102)* genomic clone into a *kin-4(null)* background can alter the thermotaxis phenotype. While introduction of the *kin-4(+)* clone rescued the thermotaxis defect of the *kin-4(tm1049)* animals, *kin-4(tm1049)* animals carrying the *kin-4(nj102)* clone displayed a thermophilic phenotype (Fig. 5E). By contrast, introduction of the *kin-4(tm1049)* genomic clone into *kin-4(nj102)* animals did not affect the thermotaxis phenotype (Fig. 5B). These results suggested that *kin-4(nj102)* is a reduction-of-function mutation.

Our results indicated that *kin-4* can be mutated to cause either a thermophilic or a cryophilic phenotype, suggesting that *kin-4* plays dual roles in regulating thermotaxis, with one activity promoting a thermophilic drive and the other a cryophilic movement. That the *kin-4* null phenotype displayed a bimodal distribution on the temperature gradient is consistent with this notion. Our previous study showed that the AFD-specific expression of a wild-type *kin-4* cDNA rescued the *kin-4* null phenotype (Nakano *et al*. 2020). These observations support that KIN-4 exerts these opposing thermophilic and cryophilic controls within the AFD neurons and suggest that *kin-4* would be a master regulator of single-cell computation by the AFD neuron.

What is the nature of the allele of *kin-4* that causes a thermophilic phenotype? Although the gene dosage analysis of *kin-4* suggested that *kin-4(nj102)* would be a reduction-of-function mutation, *kin-4(nj102)* might not be a simple reduction of function mutation, since heterozygous animals of a *kin-4* null mutation, *kin-4(tm1049Δ)*, showed the wild-type phenotype (Fig. 5D). Our observations indicated that both thermophilic alleles of *kin-4, nj102* and *nj171Δ*, are predicted to remove the PDZ domain from the KIN-4 protein. These results raise the possibility that the PDZ domain of KIN-4 is specifically engaged in the KIN-4 cryophilic activity but is dispensable for its thermophilic drive. We speculate that thermophilic alleles of *kin-4* might result from elimination of the PDZ domain, which causes the reduction specifically in the thermophilic activity of *kin-4* while maintaining the cryophilic activity. Since the PDZ domain is known to be involved in protein-protein interactions, it would be important to determine the interaction partner of KIN-4 through its PDZ domain (An *et al*. 2019). Such analysis might uncover the molecular basis underlying the dual roles of KIN-4 in the AFD thermosensory neurons for the regulation of thermotaxis.

### A genetic screen for suppressors of *mec-2(nj89gf)*

Among the thermophilic mutants we had isolated, we have previously shown that *nj89* is a gain-of-function allele of the gene *mec-2* (Nakano *et al*. 2020), which encodes a *C. elegans* homolog of stomatin (Huang *et al*. 1995). We showed that *mec-2* functions in the AFD thermosensory neurons to regulate thermotaxis and that the *mec-2(nj89gf)* mutation affected the neural activity of the AIY interneuron (Nakano *et al*. 2020), which is directly innervated by the AFD thermosensory neurons (White *et al*. 1986; Cook *et al*. 2019). However, the molecular mechanisms by which MEC-2 regulates the AIY neural activity remained elusive.

To further understand the mechanisms by which *mec-2* controls the AIY neural activity and consequently thermotaxis behavior, we conducted another genetic screen to look for mutations that can suppress the thermophilic phenotype of *mec-2(nj89gf)*. We mutagenized *mec-2(nj89gf)* animals and looked for animals that displayed cryophilic phenotypes (Fig. 6A). From this screen, we isolated 13 mutations - *nj254, nj255, nj256, nj260, nj262, nj263, nj267, nj268, nj269, nj270, nj271* and *nj274* - that altered the *mec-2(nj89gf)* phenotype. We have previously shown that *nj271* and *nj274* are alleles of *dgk-1*, which encodes a diacylglycerol kinase. *dgk-1* also functions in the AFD thermosensory neurons and affects the neuronal response of the AIY interneuron (Nakano *et al*. 2020).

**Figure 6.**
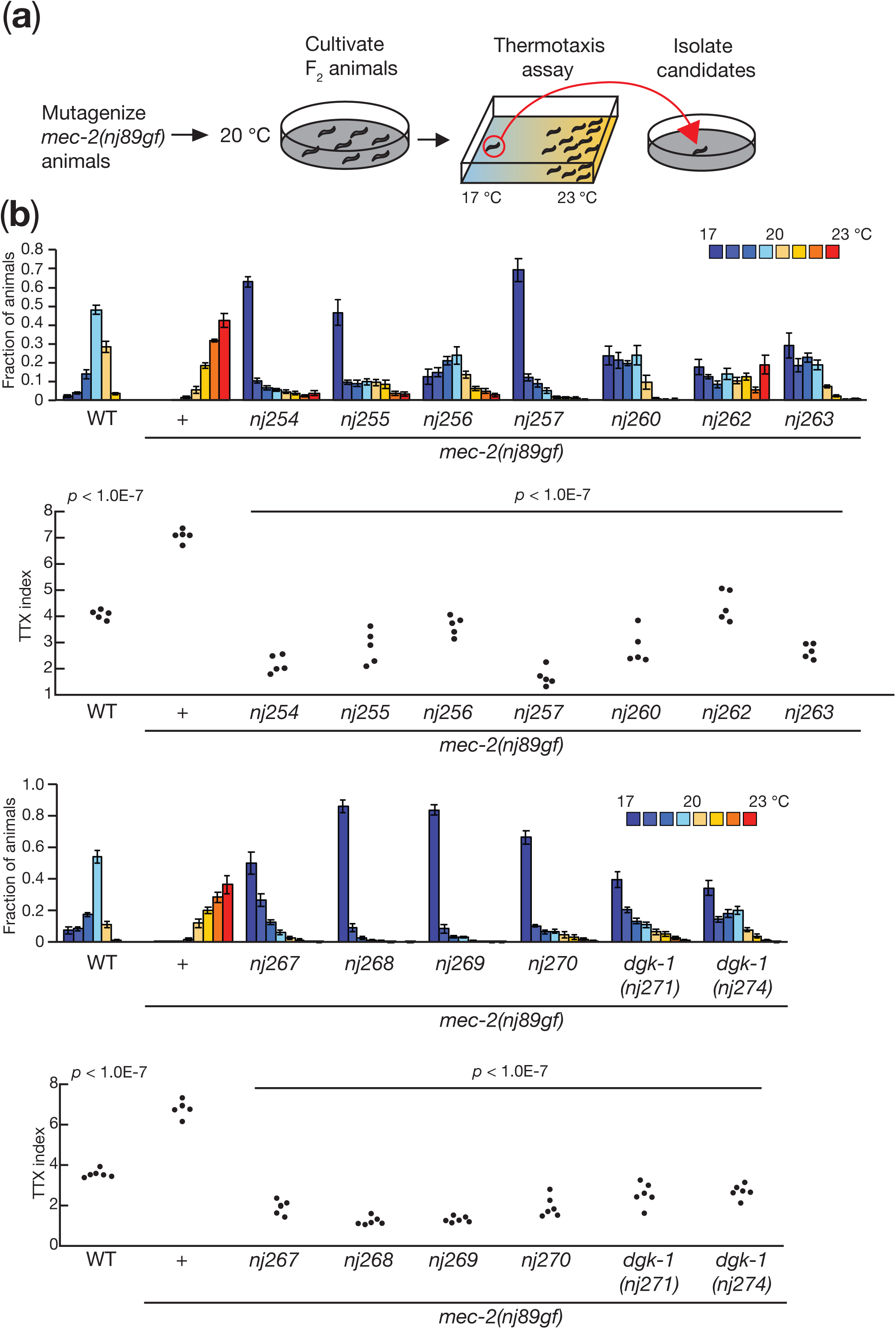
A genetic screen identified mutations that suppressed the thermophilic defect of *mec-2(nj89gf)*. a) Schematic of the genetic screen. We mutagenized *mec-2(nj89gf)* animals, and their F_2_ progeny cultivated at 20 °C were subjected to thermotaxis assays. Animals that had migrated toward the 17 °C region were isolated as mutant candidates. b)-c) Thermotaxis behaviors of mutant strains isolated from the *mec-2(nj89gf)* suppressor screen. Distributions of animals on the temperature gradients were shown as means ± SEM. TTX indices were indicated as dots. *P* values were determined by one-way ANOVA with Tukey-Kramer test. *P* values indicate the comparison of the wild-type and each suppressor isolates to *mec-2(nj89gf)*.

### *nj260* and *nj263* are alleles of *ttx-3*

Amongst the isolates we recovered from the *mec-2(nj89gf)* suppressor screen, we observed that *nj260* and *nj263* failed to complement each other. We mapped *nj260* mutation into chromosome X and found that *nj260* carries a mutation in the gene *ttx-3*, which encodes a LIM homeodomain transcription factor required for the cell fate specification of the AIY interneuron (Hobert *et al*. 1997). *nj260* animals are associated with a C-to-T transition mutation that alters the proline 371 codon of *ttx-3a* into a serine codon (Fig. 7A). We could not identify a mutation in the *ttx-3* locus of *nj263*. We attempted to amplify the *ttx-3* locus from *nj263* by polymerase chain reaction (PCR) with multiple primer sets but could not obtain PCR fragments. To assess whether *nj263* is an allele of *ttx-3*, we introduced a genomic PCR product containing the *ttx-3* locus into *mec-2(nj89gf) nj263* animals and observed that the transgenic animals at least partly reverted to the thermophilic phenotype (Fig. 7B). These results indicated that *nj260* and *nj263* are alleles of *ttx-3*. We speculate that *nj263* might carry a complex chromosomal rearrangement that involves the *ttx-3* locus. That loss of *ttx-3* function suppresses the thermophilic defect of *mec-2(nj89gf)* is consistent with our previous observation that *mec-2(nj89gf)* affects thermotaxis through the regulation of the AIY neural activity (Nakano *et al*. 2020).

**Figure 7.**
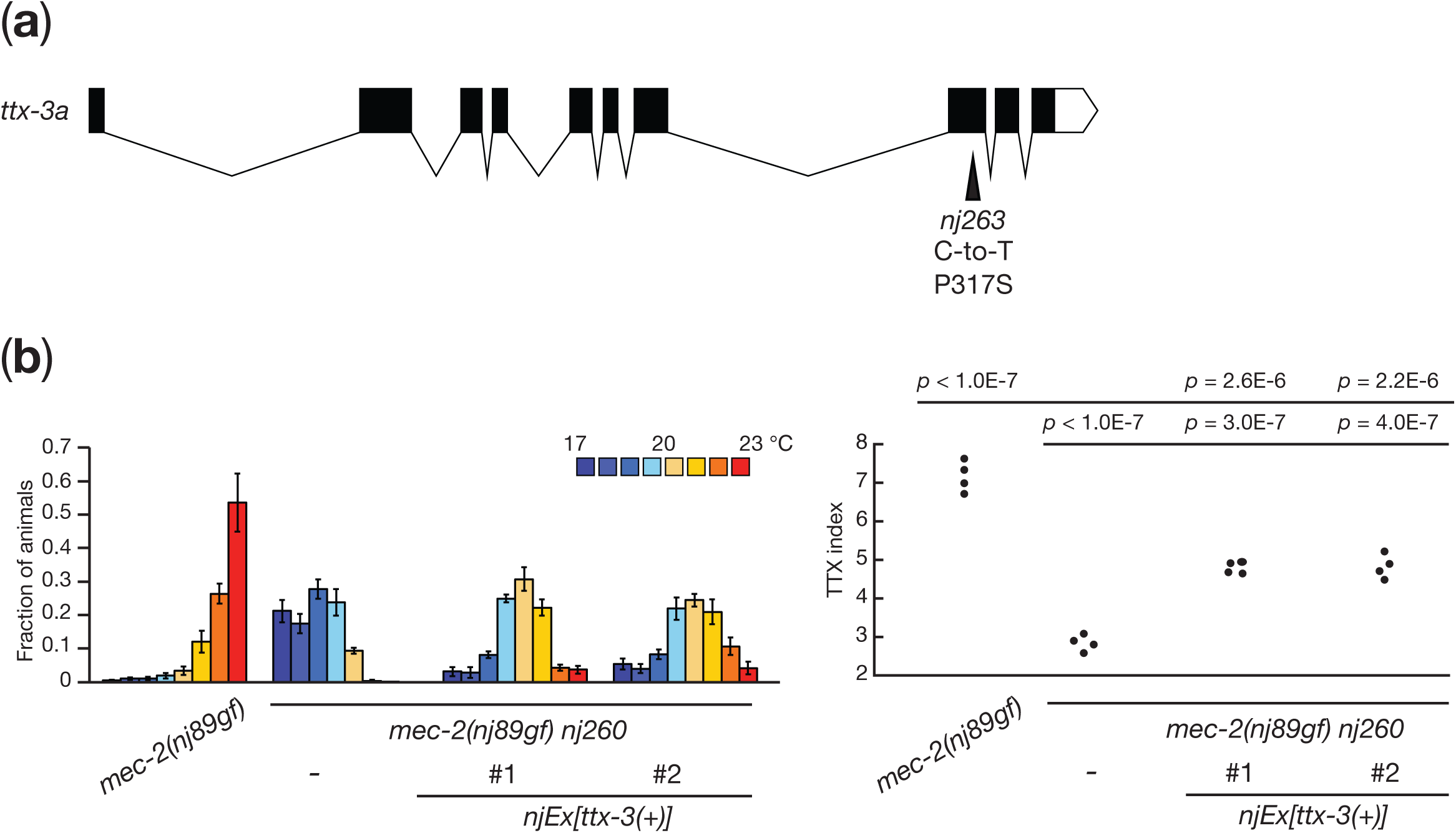
Mutations in *ttx-3* suppressed the thermophilic defect of *mec-2(nj89gf)*. a) A gene structure of *ttx-3a* and a mutation found in *nj263* are shown. The black boxes indicate exons, the white box untranslated sequence, and the lines introns. b) Thermotaxis behaviors of *mec-2(nj89gf)* and *mec-2(nj89gf) nj260* animals with or without a transgene containing a genomic fragment of the *ttx-3* locus. Distributions of animals on the temperature gradients were shown as means ± SEM. TTX indices were indicated as dots. *P* values were determined by one-way ANOVA with Tukey-Kramer test.

### Loss of *crh-1* function can suppress the thermotaxis defect of *mec-2(nj89gf)*

To identify the gene responsible for *nj257*, we first outcrossed *mec-2(nj89gf); nj257* animals and isolated *nj257* in an *mec-2(+)* background by following the activity that causes the cryophilic phenotype. Using this *nj257* mutant strain, we mapped the mutation into a 45 kb interval of chromosome III (Fig. 8A). This region contains the gene *crh-1*, which encodes a *C. elegans* homolog of CREB transcription factor (Kimura *et al*. 2002). We previously showed that *crh-1* is required for thermotaxis and that *crh-1* functions in the AFD thermosensory neurons to regulate thermotaxis (Nishida *et al*. 2011). DNA sequencing analysis of *nj257* animals identified a G-to-A transition mutation that is predicted to alter the arginine 282 codon of *crh-1a* to a histidine codon (Fig. 8A). A pan-neuronal expression of a *crh-1* cDNA using an *unc-14* promoter rescued the cryophilic defect of *nj257* animals (Fig. 8B). We also generated a deletion allele of *crh-1, nj366*, which is predicted to eliminate the entire DNA binding domain of CRH-1 and is thus likely a null allele of *crh-1*. Like *nj257, crh-1(nj366)* displayed a cryophilic phenotype (Fig. 8C). We also confirmed that *crh-1(tz2)*, another deletion allele of *crh-1* (Kimura *et al*. 2002), showed a cryophilic phenotype similar to those observed in *crh-1(nj257)* and *crh-1(nj366)* (Fig. 8C). These results established that *nj257* is an allele of *crh-1* and that loss of *crh-1* function can suppress the thermophilic defect of *mec-2(nj89gf)*.

**Figure 8.**
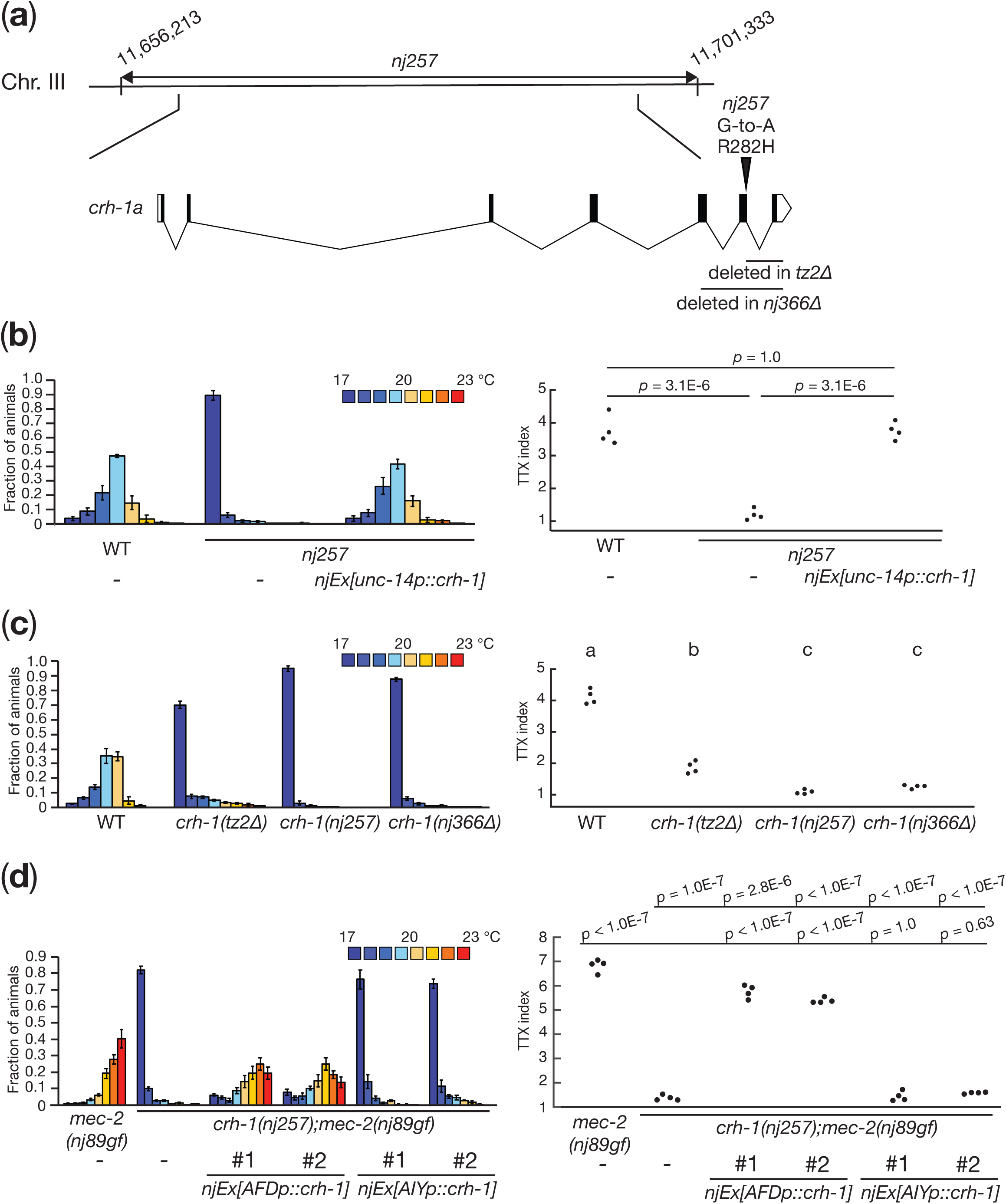
*crh-1(nj257)* suppressed the thermophilic defect of *mec-2(nj89gf)*. a) A chromosomal region to which *nj257* was mapped is indicated. The gene structure of *crh-1a* and mutations associated with each mutant are shown. The black boxes indicate exons, the lines introns, and the white boxes untranslated sequences. b)-d) Thermotaxis behaviors of *crh-1* mutants. Distributions of animals on the temperature gradients were shown as means ± SEM. TTX indices were indicated as dots. *P* values were determined by one-way ANOVA with Tukey-Kramer test.

To identify the site of *crh-1* action for the suppression of the thermophilic phenotype conferred by *mec-2(nj89gf)*, we conducted a cell-specific rescue experiment. We expressed a *crh-1* cDNA specifically in the AFD or the AIY neurons of *crh-1(nj257); mec-2(nj89gf)* animals and observed that animals expressing *crh-1* in AFD reverted to the thermophilic phenotype, while animals expressing *crh-1* in AIY did not (Fig. 8D). These results indicated that loss of *crh-1* function in AFD can suppress the thermotaxis defect of *mec-2(nj89gf)* and suggested that *crh-1* acts downstream of, or in parallel to, *mec-2* in AFD to regulate thermotaxis.

### *crh-1(nj257)* suppressed the defect of the AIY calcium response of *mec-2(nj89gf)*

We previously showed that the AIY neurons exhibit bidirectional neural responses that correlate with the valence of thermal stimuli: AIY is excited when temperature is increased toward the cultivation temperature, while AIY is inhibited when temperature is increased away from the cultivation temperature. While the temperature-evoked Ca^2+^ responses in the AFD thermosensory neurons are normal in *mec-2(nj89gf)* mutants, they showed a defect in this bidirectional AIY response. The AIY neurons of *mec-2(nj89gf)* animals displayed excitatory responses even when temperature was increased away from the cultivation temperature (Nakano *et al*. 2020).

To investigate the neural mechanisms underlying the *crh-*1-dependent suppression of *mec-2(gf)*, we conducted Ca^2+^ imaging experiments. We first examined temperature-evoked Ca^2+^ responses of the AFD thermosensory neurons in immobilized animals. The AFD neurons respond to warming stimuli by increasing the intracellular Ca^2+^ level (Kimura *et al*. 2004; Clark *et al*. 2006; Ramot *et al*. 2008; Kobayashi *et al*. 2016; Tsukada *et al*. 2016; Takeishi *et al*. 2016). We previously showed that the Ca^2+^ responses of the AFD neuron in *mec-2(nj89gf)* animals were indistinguishable from that of the wild-type animals (Nakano *et al*. 2020). When *crh-1(nj257); mec-2(nj89gf)* animals were subjected to temperature ramps, the AFD neurons increased the Ca^2+^ levels similarly to those observed in *mec-2(nj89gf)* animals (Fig. 9A). These results indicate that *crh-1* affects a process downstream of the Ca^2+^ influx in the AFD neurons to suppress the thermotaxis phenotype of *mec-2(nj89gf)*.

**Figure 9.**
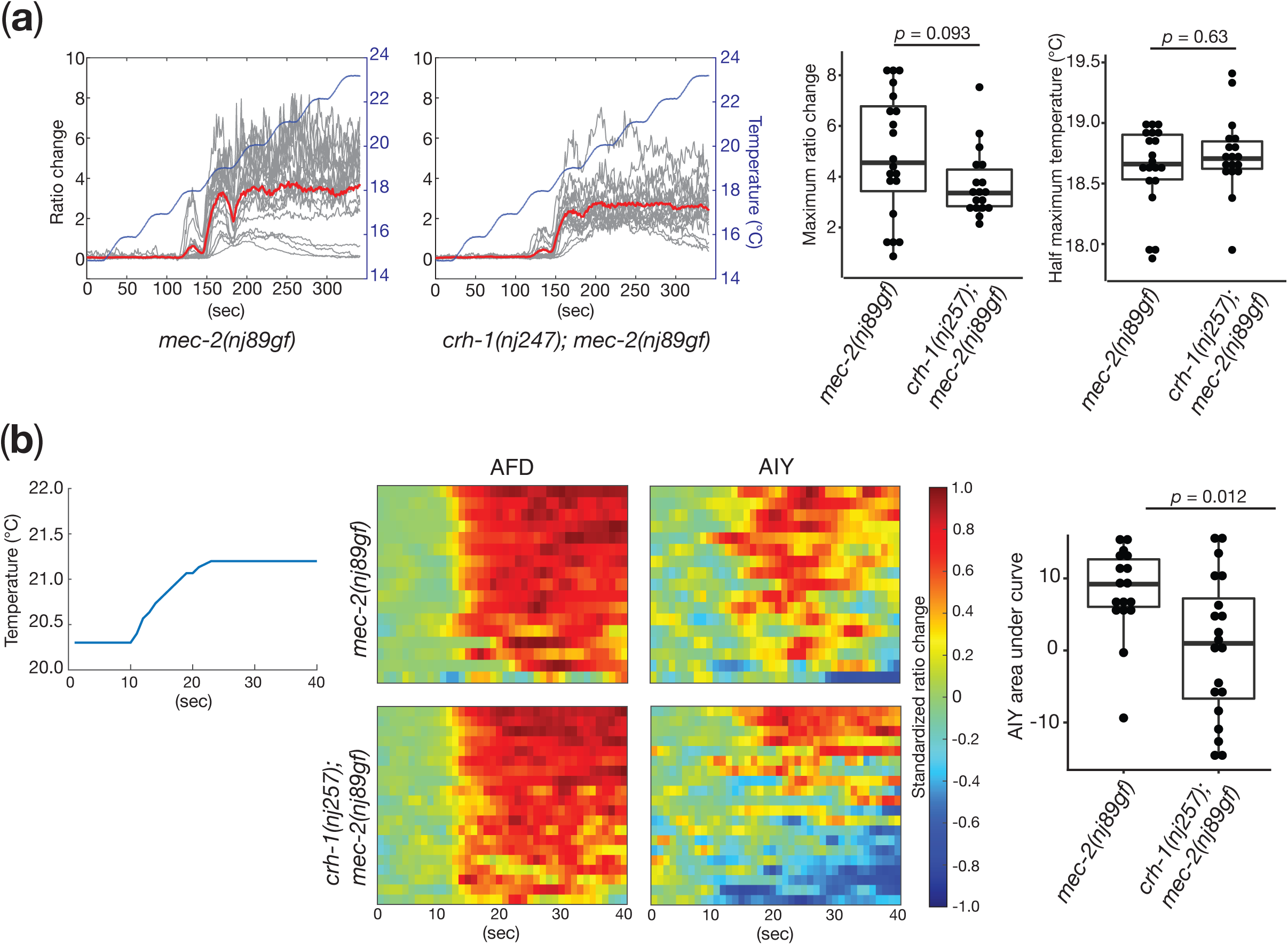
*crh-1* regulates the AIY neuronal activity. a) Calcium imaging of the AFD thermosensory neurons in immobilized animals expressing the calcium indicator, *YCX1*.*6*. The temperature stimulus is shown in the blue lines. Individual calcium responses are shown as the ratio changes of YCX/CFP in gray lines. The mean responses are indicated in the red lines. The box and dot plots of the maximum ratio change and the half maximum temperature are shown. The boxes display the first and third quartiles, the lines inside the boxes are the medians, and the whiskers extend to 1.5-time interquartile range from the boxes. *P* values were determined by Wilcoxon rank sum test. *n* = 20 and 18 for *mec-2(nj89gf)* and *crh-1(nj257); mec-2(nj89gf)* animals, respectively. b) Calcium imaging from freely-moving animals expressing *YCX1*.*6* in the AFD and AIY neurons. A representative of the temperature stimulus is shown. The heatmaps indicate the standardized ratio change of the AFD and AIY calcium dynamics. The areas under the curve of the AIY standardized ratio changes are represented in the box and dot plots. The boxes indicate display the first and third quartiles, the lines inside the boxes are the medians, and the whiskers extend to 1.5-time interquartile range from the boxes. *P* values were determined by Wilcoxon rank sum test. *n* = 17 and 20 for *mec-2(nj89gf)* and *crh-1(nj257); mec-2(nj89gf)* animals, respectively.

We next asked whether *crh-1* regulates the neuronal activity of the AIY interneuron. Since the AIY neural activity is likely influenced by the motor states of the animal (Luo *et al*. 2014; Li *et al*. 2014), we conducted the imaging from freely-moving animals. We cultivated animals at 20 °C and conducted simultaneous Ca^2+^ imaging of the AFD and AIY neurons from freely-moving animals. We subjected the animals to a temperature ramp that increases from 20.2 °C to 21.2 °C. As previously reported (Nakano *et al*. 2020), the AIY neurons of *mec-2(nj89gf)* animals predominantly exhibited excitatory responses under this condition (Fig. 9B). By contrast, a significant fraction of the AIY neurons from *crh-1(nj257); mec-2(nj89gf)* animals displayed inhibitory responses (Fig. 9B). The AFD neurons from both *mec-2(nj89gf)* and *crh-1(nj257); mec-2(nj89gf)* animals showed increases in the Ca^2+^ concentrations upon the warming stimuli. These results suggest that *crh-1* functions in the AFD neurons and regulates the bidirectional responses of the AIY interneurons to regulate thermotaxis.

Our previous study indicated that *crh-1* controls the excitability of the AFD thermosensory neurons in response to certain thermal stimuli (Nishida *et al*. 2011). Our observations indicated that in addition to this role in regulating the AFD neural activity, *crh-1* controls the neuronal outputs from AFD, thereby governing the bidirectional AIY activity. We suggest that *crh-1* might regulate transcription of a set of genes in AFD, some of which adjust the excitability of the AFD neurons while others control the AFD neuronal output to its post-synaptic neurons AIY. Our results together with our previous observations thus highlight the dual roles of *crh-1* within the AFD neurons for the regulation of thermotaxis.

Our results also suggested that *mec-2* could act upstream of *crh-1* in the AFD thermosensory neurons to regulate thermotaxis. Since *crh-1* is a transcriptional regulator, these observations raised a possibility that *mec-2* would regulate thermotaxis by controlling transcription of genes within AFD that either directly or indirectly affect the neuronal outputs from the AFD neurons. Previous studies indicated that when the cultivation temperature was shifted, the wild-type animals required certain time to adjust their thermotaxis behavior (Hedgecock and Russell 1975; Mohri *et al*. 2005; Aoki *et al*. 2018; Hawk *et al*. 2018). The adaptation to new cultivation temperature involves transcriptional reconfiguration of genes expressed in the AFD thermosensory neurons (Yu *et al*. 2014). Our previous observations also indicated that *crh-1* is required to promote the adaptation to new cultivation temperature (Nishida *et al*. 2011). These findings thus suggest that the AFD neurons in *mec-2* mutants might be defective in setting the level of gene expression appropriate for the cultivation temperature. Since the AFD neurons apparently compute its neuronal outputs based on the cultivation temperature and current thermal context (Hawk *et al*. 2018; Nakano *et al*. 2020), such a defect in AFD of *mec-2* mutants would result in an abnormal neuronal output from AFD, leading to a defect in the bidirectional AIY activity. Thus, in contrast to previous studies that indicated the roles of the stomatin family proteins in regulating the ion channels (Goodman *et al*. 2002; Price *et al*. 2004), our genetic screens suggest a new mode of the MEC-2/stomatin action that involves the transcriptional regulation in controlling the dynamics of a neural circuitry.

## Data availability

*C. elegans* strains and plasmids are available upon request. FileS1 contains descriptions of the strains used in this study. FileS2 contains numeric data of imaging analyses from freely-moving animals. The authors affirm that all data necessary for confirming the conclusions of the article are present within the article, figures and supplemental materials.

## Acknowledgements

We thank S. Mitani at National BioResource for strains; K. Ikegami, Y. Murakami and M. Murase for technical and administrative assistance; the members of the Mori and Noma laboratories for discussions. Some strains were provided by *Caenorhabditis* Genetic Center, which is funded by NIH Office of Research Infrastructure Programs (P40 OD010440).

